# *Xanthomonas* infection transforms the apoplast into an accessible and habitable niche for *Salmonella enterica*

**DOI:** 10.1101/2022.03.28.486164

**Authors:** Megan H. Dixon, Kimberly N. Cowles, Sonia C. Zaacks, Isabel N. Marciniak, Jeri D. Barak

## Abstract

The physiology of plant hosts can be dramatically altered by phytopathogens. *Xanthomonas hortorum* pv. *gardneri* is one such pathogen that creates an aqueous niche within the leaf apoplast by manipulating the plant via the transcription activator-like effector AvrHah1. Simultaneous immigration of *X. gardneri* and *Salmonella enterica* to healthy tomato leaves results in increased survival of *S. enterica* as *Xanthomonas* infection progresses. However, the fate of *S. enterica* following arrival on actively infected leaves has not been examined. We hypothesized that the water-soaking caused by *X. gardneri* could facilitate the ingression of *S. enterica* into the apoplast, and that this environment would be conducive for growth. We found that a water-soaked apoplast, abiotically or *Xanthomonas*-infected, enabled surface *S. enterica* to passively localize to the protective apoplast and facilitated migration of *S. enterica* to distal sites within the aqueous apoplast. *AvrHah1* contributed to the protection and migration of *S. enterica* early in *X. gardneri* infection. *Xanthomonas*-infected apoplasts facilitated prolonged survival and promoted *S. enterica* replication compared to healthy apoplasts. Access to an aqueous apoplast in general protects *S. enterica* from immediate exposure to irradiation whereas, the altered environment created by *Xanthomonas* infection provides growth-conducive conditions for *S. enterica*. Overall, we have characterized an ecological relationship in which host infection converts an unreachable niche to a habitable environment.

**Importance:** Bacterial spot disease caused by *Xanthomonas* species devastates tomato production worldwide. Salmonellosis outbreaks from consumption of raw produce have been linked to the arrival of *Salmonella enterica* on crop plants in the field via contaminated irrigation water. Considering that *Xanthomonas* is difficult to eradicate, it is highly likely that *S. enterica* arrives on leaves pre-colonized by *Xanthomonas* with infection underway. Our study demonstrates that infection and disease fundamentally alter the leaf, resulting in redistribution and change in abundance of a phyllosphere bacterial member. These findings contribute to our understanding of how *S. enterica* manages to persist on leaf tissue despite lacking the ability to liberate nutrients from plant cells. More broadly, this study reveals a mechanism by which physiochemical changes to a host environment imposed by a plant pathogen can convert an uninhabitable leaf environment into a hospitable niche for select epiphytic microbes.

## INTRODUCTION

Within an environment as dynamic and complex as the phyllosphere, water is one of the most critical factors for microorganism survival (1). The waxy cuticle limits the amount of moisture available to bacteria that colonize leaves. However, surface bacteria have been known to colonize particularly leaky areas of the cuticle such as leaf veins, cell junctions, and the base of glandular trichomes (2–4). Some phytopathogenic bacteria have adapted to the water scarcity issue by creating their own aqueous environment within the apoplast of leaf tissues (5–7).

Infection relies on first entering the leaf through stomata and then colonizing the extracellular internal space, or apoplast, within the leaf. *Xanthomonas hortorum* pv. *gardneri* (hereafter, *X. gardneri*), a causal agent of bacterial spot disease of tomato and pepper, is one such plant pathogen that infects leaves. Once *X. gardneri* has actively migrated to the apoplast, it can deploy its type-III secretion system to manipulate the host into leaking cellular constituents into this space (6), creating an aqueous environment. These physiochemical modifications greatly benefit *X. gardneri* but devastate its host. Bacterial spot disease often results in necrotic lesions on leaves and fruit and defoliation (8) which are detrimental to the health of the plant.

Diseased plants are hosts to a diverse community of microbes, which can be influenced by the drastic changes to their environment caused by phytopathogen infection (9–11). Microbes that are non-pathogenic to plants can benefit from disease. For example, enteric human pathogens such as *Salmonella enterica* lack enzymes that would enable the liberation of nutrients directly from plant cells (12, 13) and are typically forced to scavenge scant resources on the leaf surface. On healthy plant tissues, *S. enterica* populations decline over time (14, 15). However, plant pathogens and even insect pests that exist in agricultural fields have been shown to enhance *S. enterica* populations on plant leaves, and such organisms that can promote *S. enterica* persistence and/or replication have been coined “biomultipliers” (15–19). Bacterial pathogens including *Xanthomonas* spp. and *Pectobacterium carotovorum* have been implicated as biomultipliers for *S. enterica* on leaves (15–17, 20, 21). The mechanisms explaining how these organisms benefit *S. enterica* are not well understood, posing ecological questions with important agricultural and human disease epidemiological implications.

The prevalence of food-borne illness outbreaks linked to consumption of raw produce has climbed in the past few decades (22). Such outbreaks have been linked to pre-harvest contamination of crops in the field via irrigation water (23, 24). The presence of enteric pathogens within internal plant tissues has been a concern of food safety researchers, especially considering efforts to mitigate outbreaks through post-harvest sanitation of produce. Various studies investigated *S. enterica* colonization of the leaf apoplast, and the extent to which *S. enterica* can colonize this protective niche varies based on host and method. The flagella-dependent localization of *S. enterica* to internal tissue in response to light has been demonstrated via confocal microscopy of lettuce leaf pieces submerged in *S. enterica* inoculum (25). However, tomato leaves have been shown to be impermeable to *S. enterica* internalization when compared with lettuce leaves under comparable conditions (26, 27). Whether or not phytopathogen infection could enhance the permeability of leaves to enteric pathogens was not addressed by these studies.

*Xanthomonas* infection promotes *S. enterica* survival when both the plant and human pathogen arrive on healthy tomato leaves simultaneously (15, 16, 21), however, the fate of *S. enterica* on leaves actively infected with *Xanthomonas* spp. has not been described. *X. gardneri* infection can facilitate the migration of surface *X. gardneri* cells to distal tissue via absorption of *X. gardneri* suspensions into needle-damaged tissue (6), suggesting that water-soaking may promote the ingression of bacteria into the apoplast of leaves when external water is present. Water-soaking has been demonstrated to be enhanced by *X. gardneri* type-III effector AvrHah1 (6, 28). *AvrHah1* encodes a transcription activator-like effector that binds host DNA and upregulates the activity of a host pectate lyase (6), forcing the host to degrade the pectin “cement” that joins adjacent plant cells, resulting in an aqueous apoplast (29). Data from our lab has shown that *avrHah1* expression by *X. gardneri* is necessary for enhancing the persistence of *S. enterica* on tomato leaves when co-inoculated with *X. gardneri* (21). Based on these observations, we hypothesized that *avrHah1-*dependent water-soaking and aqueous apoplasts in general could serve as a mechanism for epiphytic *S. enterica* to enter protected sites of leaves.

The goal of our study was to examine the fate of epiphytic *S. enterica* following arrival on the surface of infected leaves. We examined whether an abiotically water-soaked or *Xanthomonas*-infected apoplast could facilitate the relocation of *S. enterica* to protected niches, particularly in the absence of leaf wounding. *S. enterica* populations were considered protected if they survived UV irradiation applied to leaves. We also identified the roles of *avrHah1*, *Xanthomonas* infection progress, and *S. enterica* motility on *S. enterica* entry into the apoplast and subsequent migration beyond the arrival site. Lastly, we investigated whether the aqueous niche created by *Xanthomonas* species enhanced the prolonged survival of *S. enterica* within the apoplast to determine whether factors aside from water availability contribute to *S. enterica* survival in the apoplast. Identifying host and microbial factors that influence the redistribution and survival of a model epiphyte will broaden our understanding of microbe-facilitated adaptations by plant-associated bacteria.

## METHODS

### Bacterial Strains, Media, and Culture Conditions

Bacterial strains used in this study are described in Table 1. *Xanthomonas* cultures were prepared by inoculating nutrient broth (NB) with frozen cells and incubating at 28°C with shaking at 200 rpm for two days. *S. enterica* cultures were prepared similarly but grown in lysogeny broth (LB) and incubated overnight. Cultures were prepared to desired cell concentrations via dilution in sterile water as described below.

**Table 1:**
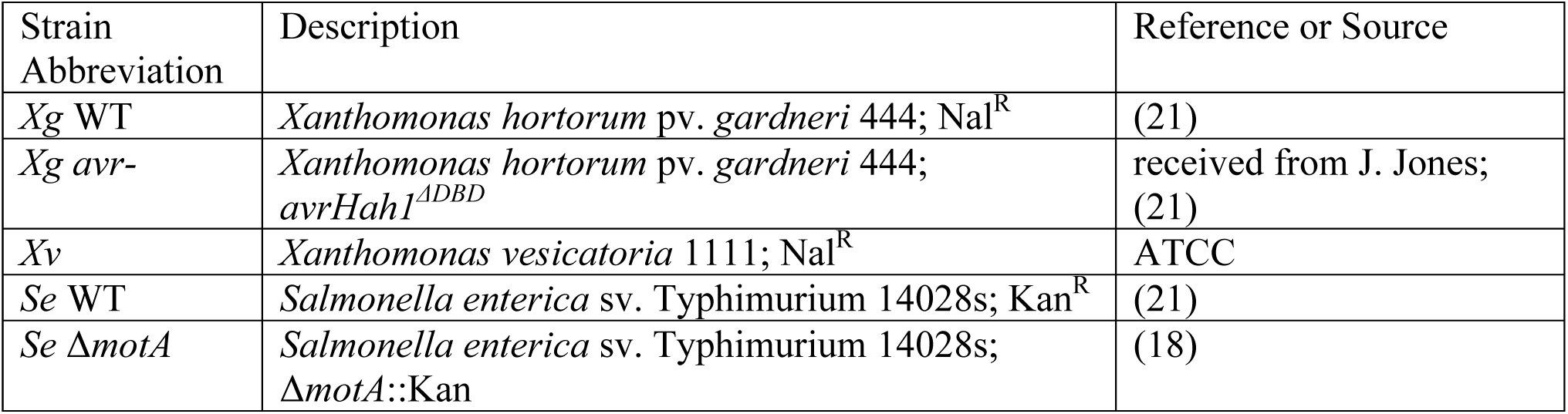
Bacterial strains used in this study.

### Plant Infiltration

For all experiments, *Solanum lycopersicum* (tomato) cultivar Moneymaker seedlings were cultivated in growth chambers with a 16 hr photoperiod at 24°C. For *Xanthomonas* infiltration experiments, 5-week-old tomato plants were infiltrated with *Xanthomonas* and inoculated with droplets of *S. enterica* as described previously (6) with modifications (Fig. 1). *Xanthomonas* cultures were prepared to an optical density at 600 nm (OD_600_) = 0.3, 0.25, or 0.2 (10^8^ CFU/mL) for *X. gardneri* WT, *X. gardneri avrHah1^ΔDBD^,* and *X. vesicatoria*, respectively, and infiltrated into two leaflets per plant on the third true leaf via needleless syringe. Infiltration medium was gently forced into the leaf tissue through the abaxial leaf surface until the infiltration zone was approximately 2 cm long, and then the infiltration zone was outlined with permanent marker.

**Figure 1:**
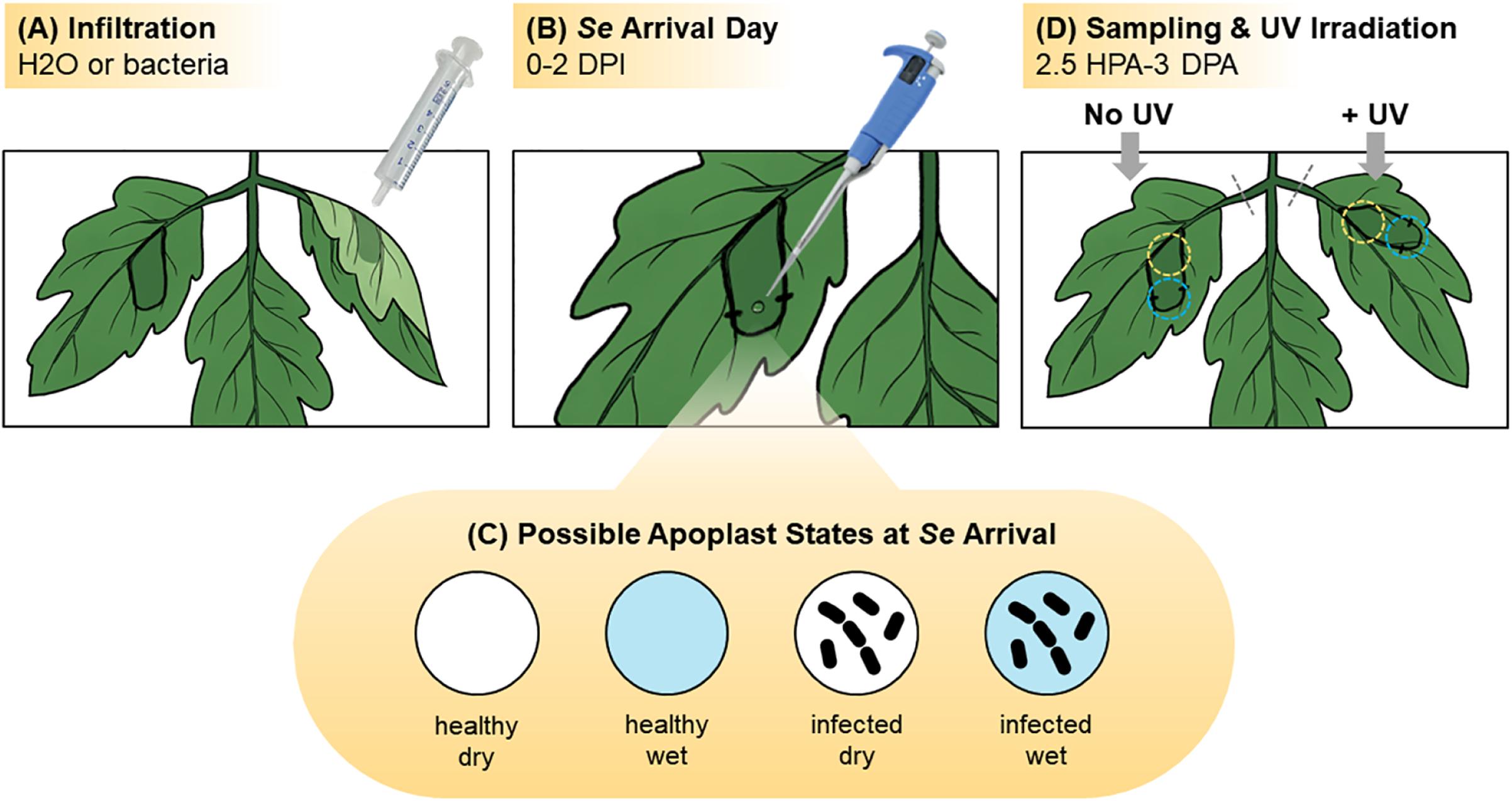
Infiltration and droplet inoculation assay. (A) Tomato leaflets were infiltrated with water or bacteria through the abaxial surface; infiltration zones are indicated with a thick black line and darker green shading. (B) Droplets of *S. enterica* (*Se*) cell suspension were then applied to the adaxial surface of each infiltration zone at a site marked with permanent marker, represented by black dash marks. The *Se* arrival day occurred 0-2 days post-infiltration (DPI). (C) When the *Se* cells arrived on the plant via droplet application, several apoplast states were possible depending on the infiltration treatment received. Circle symbols (healthy: open circle, infected: bacteria-filled circle; dry: white circle, wet: turquoise circle) depict the various apoplast states in subsequent data figures. (D) Designated leaflets were detached and irradiated with UV at indicated timepoints, which ranged from 2.5 hours post-*Se* arrival (HPA) to 3 days post-Se arrival (DPA) depending on the experiment. Tissue was sampled immediately after UV irradiation at the site of *S. enterica* arrival (blue dotted circle) and non-overlapping distal tissue (yellow dotted circles) for *Se* population enumeration.

When applicable, a subset of plants was infiltrated with sterile water as a control. When *S. enterica* infiltration was utilized as a control for one experiment, similar methods were applied, although the *S. enterica* culture used for infiltration was prepared to a concentration of 10^7^ CFU/mL and infiltrations were performed in a biosafety cabinet.

Following infiltration, plants were then incubated in bins with lids on at a relative humidity (RH) > 90% for 1-2 days prior to subsequent *S. enterica* application. For some experiments, infiltration with sterile water was delayed for a subset of plants until immediately prior to *S. enterica* inoculation to simulate arrival of *S. enterica* on healthy water-soaked tissue. These plants were incubated alongside the already infiltrated plants prior to their infiltration such that environmental conditions would be consistent among all plants within an experiment.

### Droplet Inoculation

Lids of plant bins were removed to inoculate leaflets with *S.* enterica, effectively reducing the RH to ∼70%. Infiltrated leaflets were marked again with permanent marker on the *S. enterica* arrival day to designate the site of *S. enterica* droplet placement on the infiltration zone. *S. enterica* cultures were prepared to an OD_600_= 0.1 (10^8^ CFU/mL) and 15 μL droplets of the normalized *S. enterica* culture were applied to the adaxial surface within the part of the infiltration zone closest to the leaf margin. For one experiment in which *S. enterica* was applied at a lower inoculation level, the normalized culture was diluted 1000-fold prior to inoculation. Note that droplets were applied such that they did not overlap with any minor tissue damage caused by the needleless syringe infiltration from the abaxial surface. Plants inoculated with *S. enterica* remained inside plastic bins, and the lids were kept off (RH ∼70%) until droplets were no longer visible (i.e., absorbed or evaporated). Visual observations of droplet absorption (complete, partial, or none) were recorded at 2.5 hours after droplet application. When specified, a micropipette was used for removing each remaining droplet and measuring their volume. Leaf tissue was then sampled destructively at the indicated time post-*S. enterica* arrival to quantify bacterial populations. For experiments requiring sampling beyond 2.5 hours post-*S. enterica* arrival, bins lids were replaced to maintain plants at >90% RH until sampling.

### Bacterial Population Sampling

Whole leaflets were removed at the petiolule and when specified, treated with 254 nm UV radiation (Stratalinker® UV Crosslinker 1800) at 600,000 μJ/cm^2^ with the adaxial leaf surface facing upwards towards the bulb in order to inactivate surface *S. enterica* cells and enrich for apoplastic *S. enterica*. The effect of 600,000 μJ/cm^2^ of UV irradiation on surface *S. enterica* populations was determined by removing surface *S. enterica* cells 1 day post-droplet inoculation using a cotton swab dipped in 0.1% Tween-20 and gently wiping it across the adaxial surface of the leaf for 5 seconds. The tip of the cotton swab was vortexed in 500 uL of water for 5 seconds, and then the cell mixture was dilution-plated to enumerate surface *S. enterica* populations (Fig. S1).

Leaflets irradiated with UV were chosen at random between the two leaflets sampled per plant and the second leaflet was left non-irradiated as a control. Bacterial populations were measured as described previously (18). Unless noted otherwise, all inoculated leaflets were sampled regardless of observations made at the droplet absorption stage. Briefly, leaf tissue was excised using a sterile 1 cm diameter tissue corer and homogenized in 500 μL of sterile water using a hand Dremel. For *S. enterica* droplet-inoculated tissue, the site of *S. enterica* arrival was sampled by centering the tissue corer between the marker guidelines. When indicated, tissue was also sampled distal from the arrival site by excising a second tissue core 1-2 mm away from the first leaf disk overlapping the infiltration zone. Leaf tissue homogenate was dilution-plated on LB agar amended with 50 μg/mL kanamycin for enumeration of *S. enterica* populations. When *S. enterica* was applied at a low dosage, the leaf tissue homogenate that remained after plating was enriched with LB amended with kanamycin (50 ug/mL) to determine if *S. enterica* was present, but below the limit of detection for dilution-plating. LB-enriched samples were plated 22-24 hours later. *S. enterica* colonies were enumerated after 1 day of incubation at 37°C.

### Statistical Analyses

R Studio version 2021.09.1 (R Development Core Team, R Foundation for Statistical Computing, Vienna, Austria [http://www.R-project.org]) was utilized for all statistical analyses and plot generation. Each experiment was repeated three times, and each boxplot represents the aggregation of three independent experiments found to be statistically similar (*P* > 0.05), unless noted otherwise in the figure legend. Within each experiment, the number of biological replicates *N* = 4-8 leaflets. For comparing *S. enterica* populations among treatment groups at the arrival tissue site, the ANOVA and post-hoc Tukey tests were used. When two leaflets were inoculated per plant, the data were fit to a linear mixed effects model using the lme4 package in which individual plants were modeled as a random effect. *S. enterica* population data at distal tissue sites were generally not found to be normally distributed due to zeroes resulting from the absence of *S. enterica* migration beyond the initial site of arrival. Therefore, for comparing *S. enterica* populations among treatment groups at the distal tissue site, the non-parametric Kruskal-Wallis and post-hoc Wilcox tests were used. In addition, the proportion of samples with detectable bacterial populations at the distal tissue site were compared using the Fisher’s Exact Test to compare the incidence of *S. enterica* colonization of distal tissue. For the low *S. enterica* dosage experiment, data from the healthy/dry treatment was excluded from the mixed effects model because the value of nearly every data point was zero. Preliminary analysis indicated that for the remaining two treatments (healthy/wet and *X.* gardneri-infected), all means were greater than zero based on one-way t-tests (*P* < 0.05). When applicable, the limit of detection minus one (i.e., 49 CFU/leaf disk) was substituted for non-detects. The significance threshold for all comparisons was *P* < 0.05.

## RESULTS

### Abiotically water-soaked apoplasts enable *S. enterica* to migrate via stomata to a UV-protected niche

To characterize the role of water in the apoplast in the migration of *S. enterica* to a protected niche and migration to sites distal to the arrival site, we developed a method involving infiltration of intact tomato leaflets, application of droplets of *S. enterica* cell suspensions, and measurement of UV-protected *S. enterica* populations (Fig. 1). Our methods were inspired by work previously published by Schwartz et al. (6), with a few key differences that included applying bacterial droplets to intact (not detached) leaflets onto the adaxial surface, refraining from wounding the leaves at the droplet application site, and utilizing UV irradiation to enrich for apoplastic *S. enterica* populations. We found that our UV irradiation methods reduced the *S. enterica* populations on the leaf surface by 2.9 logs (Fig. S1). Additionally, we manipulated the timing of *S. enterica* arrival on water-infiltrated leaves to observe how the presence of water in the apoplast influences the relocation of *S. enterica*.

Infiltration of water into the apoplast simulated abiotic water-soaking that may occur during periods of high precipitation (30). Under the tested conditions (∼70% RH), water infiltration created an obvious darkening or “wet” appearance to the tissue that lasted for a few hours (data not shown). *S. enterica* droplets were applied to two treatments: leaves with a (i) “healthy/dry” apoplast where the tissue had become macroscopically dry at time of *S. enterica* application at 2 day post-infiltration (DPI) or (ii) “healthy/wet” apoplast where tissue was still visibly dark and wet in appearance when *S. enterica* was applied within 10 minutes of water infiltration (0 DPI). A third treatment was included in which we infiltrated *S. enterica* directly into the apoplast to determine the relative amount of UV protection provided to apoplastic *S. enterica* cells.

Within a few hours of *S. enterica* arrival on the leaf, we observed the disappearance of the droplets specifically into healthy/wet tissue in the absence of any applied epidermal wounding, suggesting that the *S. enterica* suspension had been absorbed through stomata. Leaflets were observed for a duration of 2.5 hours post-application of *S. enterica,* and absorption of the entire 15 μL suspension could be observed within a minimum of 30 minutes. The success rate for complete absorption of the droplet for healthy/wet tissue varied between 50-83% among three experimental replicates, although droplets that failed to be completely absorbed within the 2.5 hour timeframe had visibly reduced in size, suggesting that they were partially absorbed. By contrast, healthy/dry leaflets had an absorption rate of 0%. Beyond the observation period, after several hours had passed, all droplets disappeared either by absorption, evaporation, or both. *S. enterica* populations were enumerated at 1 day post-arrival, and among the three treatments, UV irradiation resulted in the lowest *S. enterica* populations when *S. enterica* was applied to leaflets with a healthy/dry apoplast (Fig. 2). These surface-limited populations were reduced ∼2.5 logs by UV irradiation. Arrival on healthy/wet tissue and direct infiltration into the apoplast resulted in equivalent UV-protected populations and relatively small population reductions of ∼0.9 and ∼0.7 logs, respectively, in response to UV irradiation (Fig. 2). The increased UV protection conferred to *S. enterica* when applied to healthy water-soaked tissue suggests that an aqueous apoplast provides a gateway for surface bacteria to move inside the leaf.

**Figure 2:**
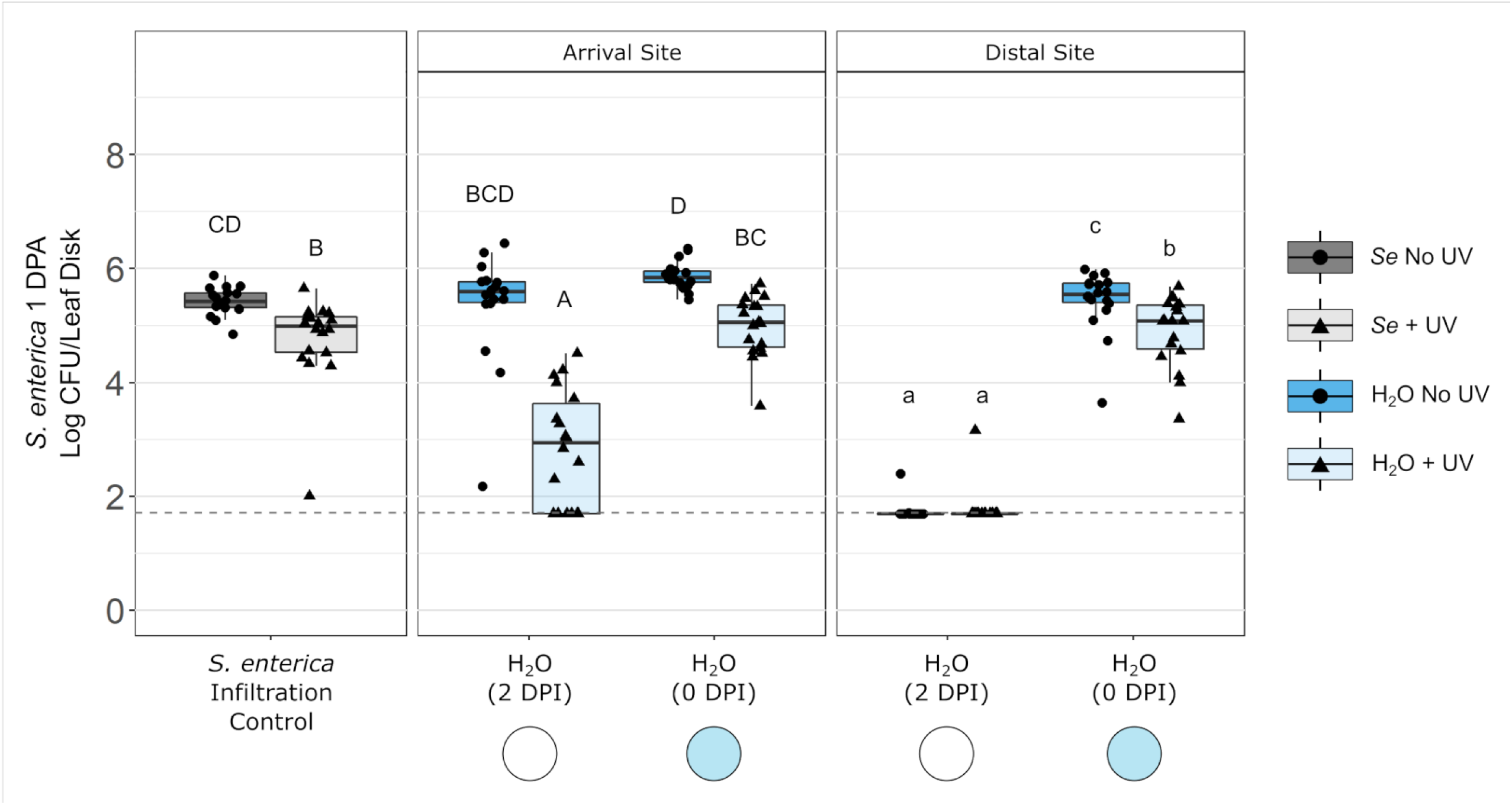
Healthy leaf tissue with a water-filled apoplast facilitates localization to a protected niche and migration to distal tissue. *S. enterica* (*Se*) was applied by droplet (10^6^ CFU/leaflet) at 2 or 0 days post-infiltration (DPI) with water (blue boxplots). Half of leaflets were UV-irradiated (triangle points; light-shaded boxplots) and the other half were left non-irradiated (circle points; dark-shaded boxplots). *Se* populations were sampled at 1 day post *Se*-arrival (DPA) at the droplet arrival site and distal from the arrival site. *Se* populations from droplet-inoculated leaflets were compared to those from control leaflets infiltrated with *S. enterica* (gray boxplots). Uppercase letters above the boxplots denote significance among treatment groups within the arrival site based on an ANOVA and post-hoc Tukey test (*P* < 0.05). Lowercase letters denote significance among treatment groups within the distal site based on a Kruskal-Wallis and post-hoc Wilcox test (*P* < 0.05). Dotted line indicates limit of detection. Refer to Figure 1, panel C for circle symbol key describing apoplast states.

Next, we investigated whether movement of surface bacteria to the apoplast was limited to the immediate substomatal chamber or extended beyond. We observed distinct differences in the ability of *S. enterica* to migrate from the arrival site to tissue non-overlapping the droplet application site depending on the physical state of the apoplast. A healthy/dry apoplast did not readily facilitate lateral migration of *S. enterica* to areas within the apoplast distal from the arrival site whereas, a healthy/wet apoplast enabled recovery of significantly higher *S. enterica* populations at distal sites (Fig. 2). Distal sites within the apoplast provided *S. enterica* with a protected niche if the apoplast was an aqueous environment at time of *S. enterica* arrival, as the population reduction in response to UV irradiation was only ∼0.5 logs (Fig. 2). These findings support the conclusion that an aqueous apoplast permits surface bacteria to migrate beyond the initial arrival site.

### *Xanthomonas* water-soaking promotes *S. enterica* entry to the apoplast and facilitates migration to distal tissue

*X. gardneri* is known to cause water-soaking (6, 28), and thus we examined whether *X. gardneri* infection could confer protection to *S. enterica* against UV irradiation and facilitate its migration to infected sites distal from the initial site similarly to a healthy/wet apoplast. We found that *X. gardneri-*infected tissue enabled complete absorption of *S. enterica* droplets into the leaves at 1 DPI (Fig. S2) and 2 DPI (images not shown) within 0.5-2.5 hours. To assess the protection of absorbed *S. enterica* cells against UV, we infiltrated an excess number of plants with *X. gardneri*, and the first 50% of plants that completely absorbed the droplets were specifically selected for *S. enterica* population sampling at 1 day post-*S. enterica* arrival. *S. enterica* cells were found to be more protected against UV irradiation following droplet absorption into *X. gardneri*-infected tissue at 2 DPI compared to *S. enterica* cells that were limited to the surface of healthy/dry tissue (Fig. 3). Like a healthy/wet apoplast, a *X. gardneri-*infected apoplast facilitated robust colonization of *S. enterica* cells to sites in the apoplast beyond the initial arrival site, where *S. enterica* was protected from UV irradiation (Fig. 3b).

**Figure 3:**
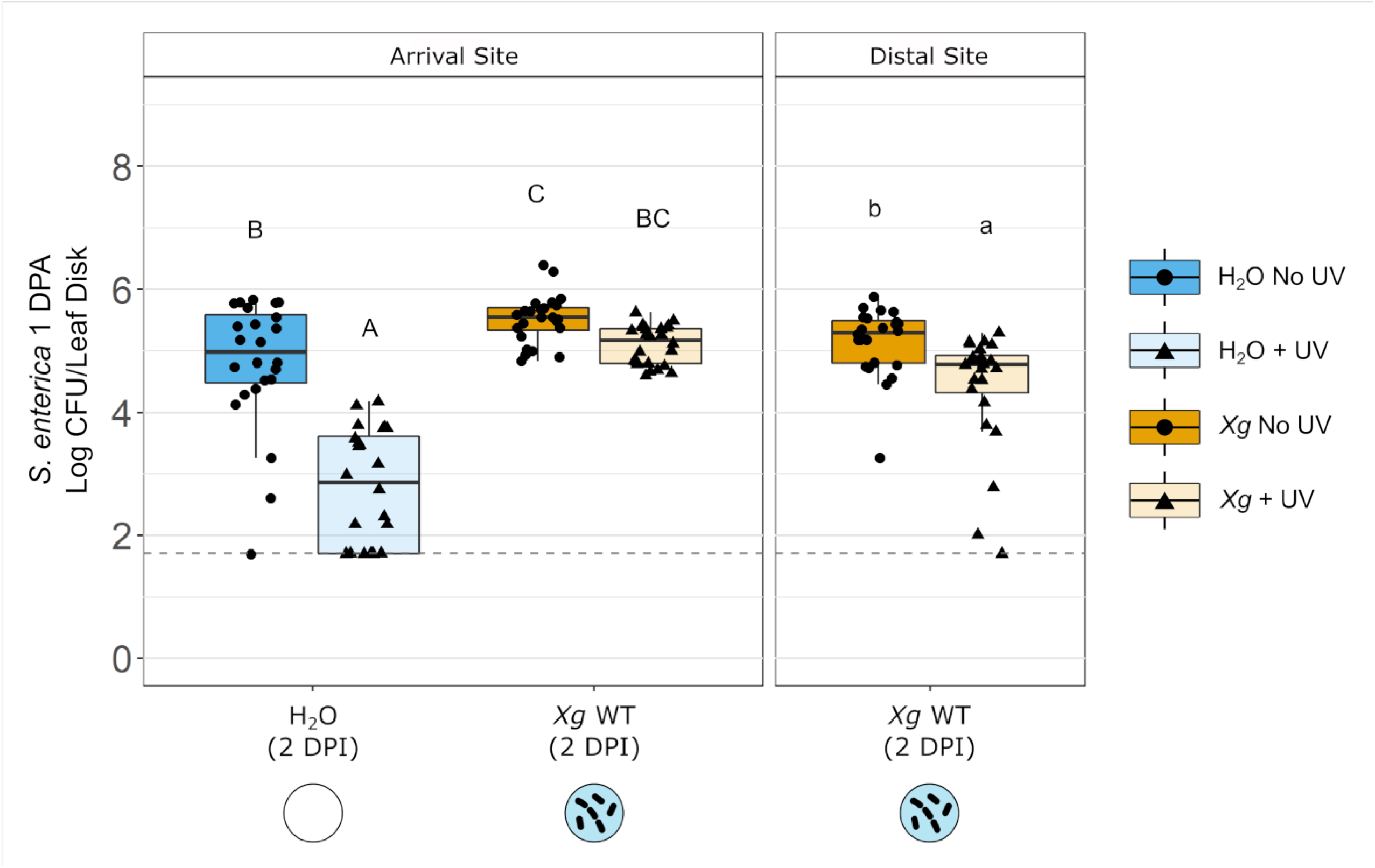
*X. gardneri* infection facilitates *S. enterica* localization to protected niche and migration beyond site of arrival. *S. enterica* (*Se*) was applied by droplet (10^6^ CFU/leaflet) at 0 days post-infiltration (DPI) with water (blue boxplots) or 2 DPI with *X. gardneri* (*Xg* WT; orange boxplots). Half of leaflets were UV-irradiated (triangle points; light-shaded boxplots) and the other half were left non-irradiated (circle points; dark-shaded boxplots). *Se* populations were sampled at 1 day post *Se*-arrival (DPA) at the droplet arrival site and distal from the arrival site (distal site for *Xg*). Uppercase letters above the boxplots denote significance among treatment groups within the arrival site based on an ANOVA and post-hoc Tukey test (*P* < 0.05). Lowercase letters denote significance among treatment groups within the distal site based on a Kruskal-Wallis and post-hoc Wilcox test (*P* < 0.05). Dotted line indicates limit of detection. Refer to Figure 1, panel C for circle symbol key describing apoplast states.

### *AvrHah1* facilitates *S. enterica* localization to the apoplast and lateral migration early in infection

Knowing that an aqueous apoplast facilitates the migration of *S. enterica* to a protected niche within a day of arrival, we examined whether protection against UV irradiation exists as early as 2.5 hours post-arrival, a time point at which *S. enterica* droplet suspensions were often completely absorbed. Also, we compared the fate of *S. enterica* following arrival on tissue infected by one of three *Xanthomonas* strains including *X. gardneri* WT, *X. gardneri avrHah1^ΔDBD^*, and *X. vesicatoria*. *X. vesicatoria* is a pathogen within the bacterial spot disease complex that lacks *avrHah1* and has been shown in co-inoculation experiments with *S. enterica* to not promote *S. enterica* growth on tomato leaves, in contrast to *X. gardneri,* which does facilitate *S. enterica* replication (15, 21).

The duration of infection with *Xanthomonas* spp. was found to affect visible water-soaking, absorption of the *S. enterica* suspension, and *S. enterica* protection from UV irradiation (Fig. 4). Tissue infected with *Xanthomonas* spp. was generally water-soaked at 1 and 2 DPI, except for leaflets infected with the *X. gardneri avrHah1^ΔDBD^* strain at 1 DPI (Fig. S3). Leaflets infected with *X. vesicatoria* became water-soaked at a similar rate as *X. gardneri* WT-infected leaflets despite lacking the *avrHah1* gene. As for effects on *S. enterica*, tissue that was macroscopically dry at *S. enterica* arrival, which included the healthy/dry and *X. gardneri avrHah1^ΔDBD^* 1 DPI treatments, did not absorb the *S. enterica* suspension, resulting in the highest droplet recovery volumes at 2.5 hours post *S. enterica* droplet application (Fig. 4a). In contrast to dry tissue, the recovered droplet volumes were lower for water-soaked tissue with healthy/wet, *X. gardneri* WT-infected, or *X. vesicatoria*-infected leaves, and these tissues often absorbed the *S. enterica* suspension completely (Fig. 4a). Congruently, *S. enterica* arrival on water-soaked tissues resulted in the greatest protection, as evident by smaller reductions in population in response to UV (Fig. 4b). Overall, infection progress had a significant effect on *S. enterica* protection in our model.

**Figure 4:**
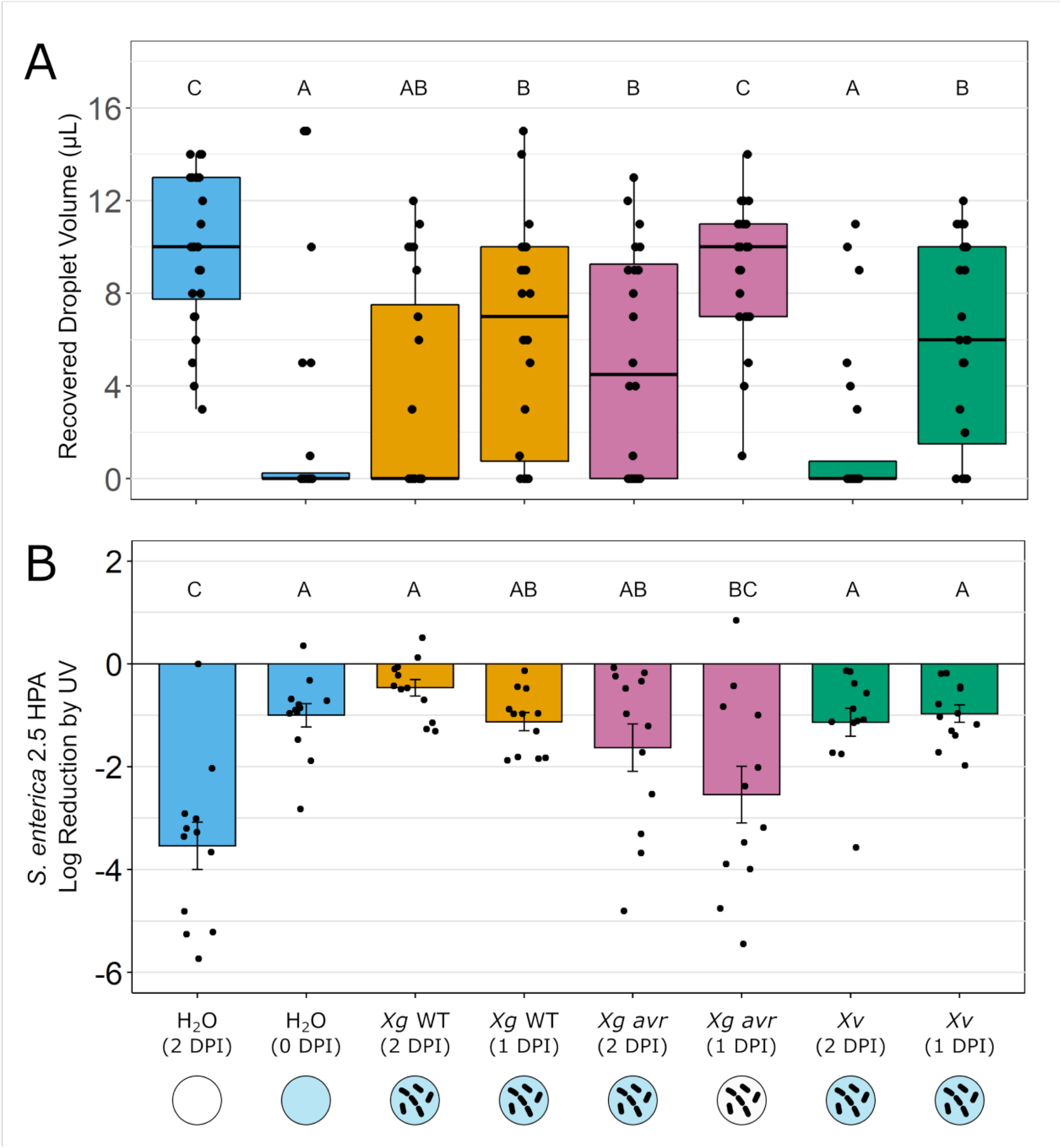
*S. enterica* access to protected niche within 2.5 hours of arrival on water-soaked tissue depends on *Xanthomonas* strain and duration of infection. *S. enterica* (*Se*) was applied by droplet (10^6^ CFU/leaflet) to leaflets infiltrated with water (blue boxplots), *X. gardneri* WT (*Xg* WT; orange boxplots), *X. gardneri avrHah1^ΔDBD^* (*Xg avr;* pink boxplots), or *X. vesicatoria* (*Xv*; green boxplots) at indicated days post-infiltration (DPI). (A) Unabsorbed *S. enterica* (*Se*) droplet volumes measured 2.5 hours following droplet application. Differing letters above the plot denote significantly different treatment groups based on a Kruskal-Wallis and post-hoc Wilcox multiple comparison test (*P* < 0.05). (B) Reduction of *S. enterica* populations in response to UV irradiation at the *Se* droplet arrival site at 2.5 hours-post *Se* arrival (HPA). Letters above the boxplots denote significance among treatment groups based on an ANOVA and post-hoc Tukey test (*P* < 0.05). Refer to Figure 1, panel C for circle symbol key describing apoplast states.

In addition to facilitating initial entry of *S. enterica* into the substomatal chamber, *avrHah1* was also found to contribute to the ability of *S. enterica* to migrate to distal tissue beyond the site of arrival early in infection (Fig. 5). Leaves infiltrated with *X. gardneri avrHah1^ΔDBD^*, which generally had a dry apoplast at 1 DPI (Fig. S3) and did not result in visible absorption of the *S. enterica* suspension (Fig. 4a), exhibited reduced migration of *S. enterica* to distal tissue compared to leaves infiltrated with *X. gardneri* WT (Fig. 5). However, by 2 DPI, when the tissue was visibly water-soaked (Fig. S3), *X. gardneri avrHah1^ΔDBD^-*infected tissue did facilitate colonization of distal tissue by *S. enterica* (Fig. 5). These findings demonstrate that the role of *avrHah1* in altering the apoplast into an aqueous environment occurs early in the disease progress, and these changes facilitate surface bacteria entry into the apoplast and subsequent lateral migration to other water-soaked areas. We also note that additional bacterial factors besides *avrHah1* contribute to water-soaking later in disease progression.

**Figure 5:**
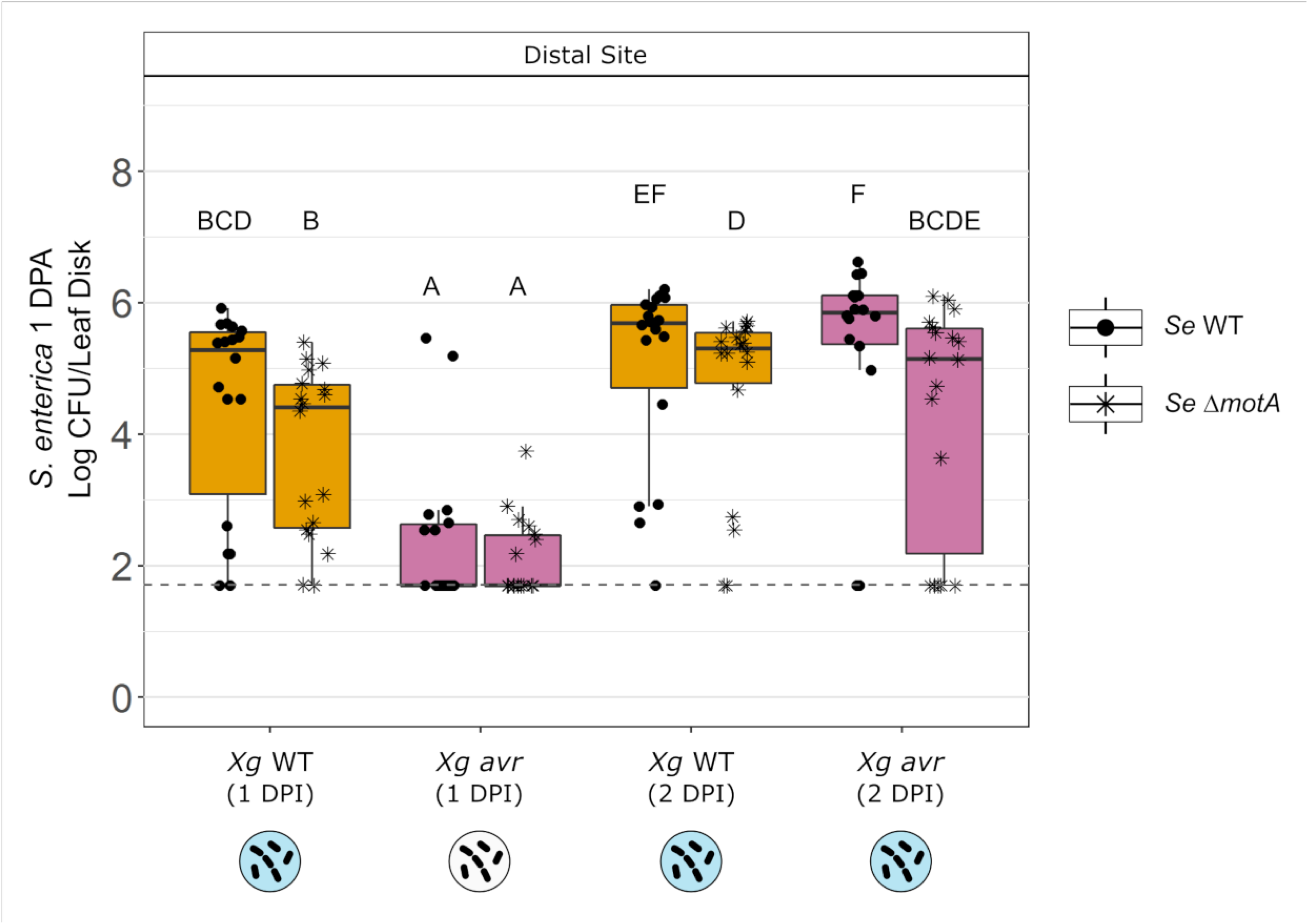
*X. gardneri avrHah1* enhances *S. enterica* migration during early infection, and flagellar motility promotes *S. enterica* survival at distal tissue. *S. enterica* (*Se*) WT (circle points) or Δ*motA* (asterisk points) was applied by droplet (10^6^ CFU/leaflet) at 1 or 2 days post- infiltration (DPI) with *X. gardneri* WT (*Xg* WT, orange boxplots) or *X. gardneri avrHah1^ΔDBD^* (*Xg avr*; pink boxplots). *Se* populations were sampled at 1 day post *Se*-arrival (DPA) distal from the *Se* droplet arrival site without UV irradiation treatment. Letters above the boxplots denote significance among treatment groups based on a Kruskal-Wallis and post-hoc Wilcox test (*P* < 0.05). Dotted line indicates limit of detection. Refer to Figure 1, panel C for circle symbol key describing apoplast states.

### Active motility of *S. enterica* increases colonization of tissue beyond the arrival site as *X. gardneri* infection progresses

To test the hypothesis that active *S. enterica* motility contributes to migration away from the surface, bacterial populations for a *S. enterica* Δ*motA* mutant with non-functional flagella were compared to a wild-type *S. enterica* strain. At 2.5 hours post-*S. enterica* arrival, the Δ*motA* mutant colonized UV-protected sites of water-soaked leaves equivalently to the wild-type *S. enterica* strain (Fig. S4), indicating that motility does not affect *S. enterica’s* ability to enter the aqueous apoplast. We also compared *S. enterica* WT and Δ*motA* populations at tissue distal from the arrival site at 1 day post-*S. enterica* arrival. Motility did not affect *S. enterica* colonization of distal tissue infected with *X. gardneri* when *S. enterica* arrived at 1 DPI, however, at 2 DPI, *S. enterica* populations at distal sites were enhanced by motility (Fig. 5). Deletion of *motA* did not eliminate *S. enterica* colonization of distal tissue nor affect the proportion of distal sites with detectable *S. enterica* populations, indicating that motility was useful but not required for lateral migration of *S. enterica*. Overall, motility was not required for entry into the apoplast or migration beyond the initial arrival site, supporting a model of passive, rather than active, *S. enterica* relocation within water-soaked tissue.

### Prolonged survival of *S. enterica* in a protected niche is increased by *Xanthomonas* infection

Having established that aqueous apoplasts result in similar migration of *S. enterica* to UV-protected niches regardless of whether the apoplast is abiotically water-soaked or water-soaked via *Xanthomonas* infection, we then tested the hypothesis that the apoplast of *Xanthomonas* infected leaves would support higher *S. enterica* population growth than healthy leaves following *S. enterica* application at 1 DPI. We found that *S. enterica* arrival on *X. gardneri* WT or *X. vesicatoria-*infected tissue resulted in the highest UV-protected *S. enterica* populations at three days post-arrival (Fig. 6). Although a healthy/wet apoplast facilitated equal colonization of *S. enterica* within UV-protected areas compared to *X. gardneri* WT and *X. vesicatoria* infected tissue at 2.5 hours post-*S. enterica* arrival, (Fig. 4b) the healthy/wet apoplast supported lower *S. enterica* populations at the arrival and distal sites relative to infected/wet apoplasts by 3 days post-*S. enterica* arrival (Fig. 6). *X. gardneri avrHah1^ΔDBD^*-infected plants (1 DPI) did not permit droplet absorption (Table S1), yielded intermediate UV-protected *S. enterica* populations at the arrival and distal sites (Fig. 6), and resulted in the most variable *S. enterica* populations by 3 DPA (Fig. 6). Healthy/dry tissue also did not absorb the droplets (Table S1) and supported the lowest *S. enterica* populations at the arrival and distal sites (Fig 6).

**Figure 6:**
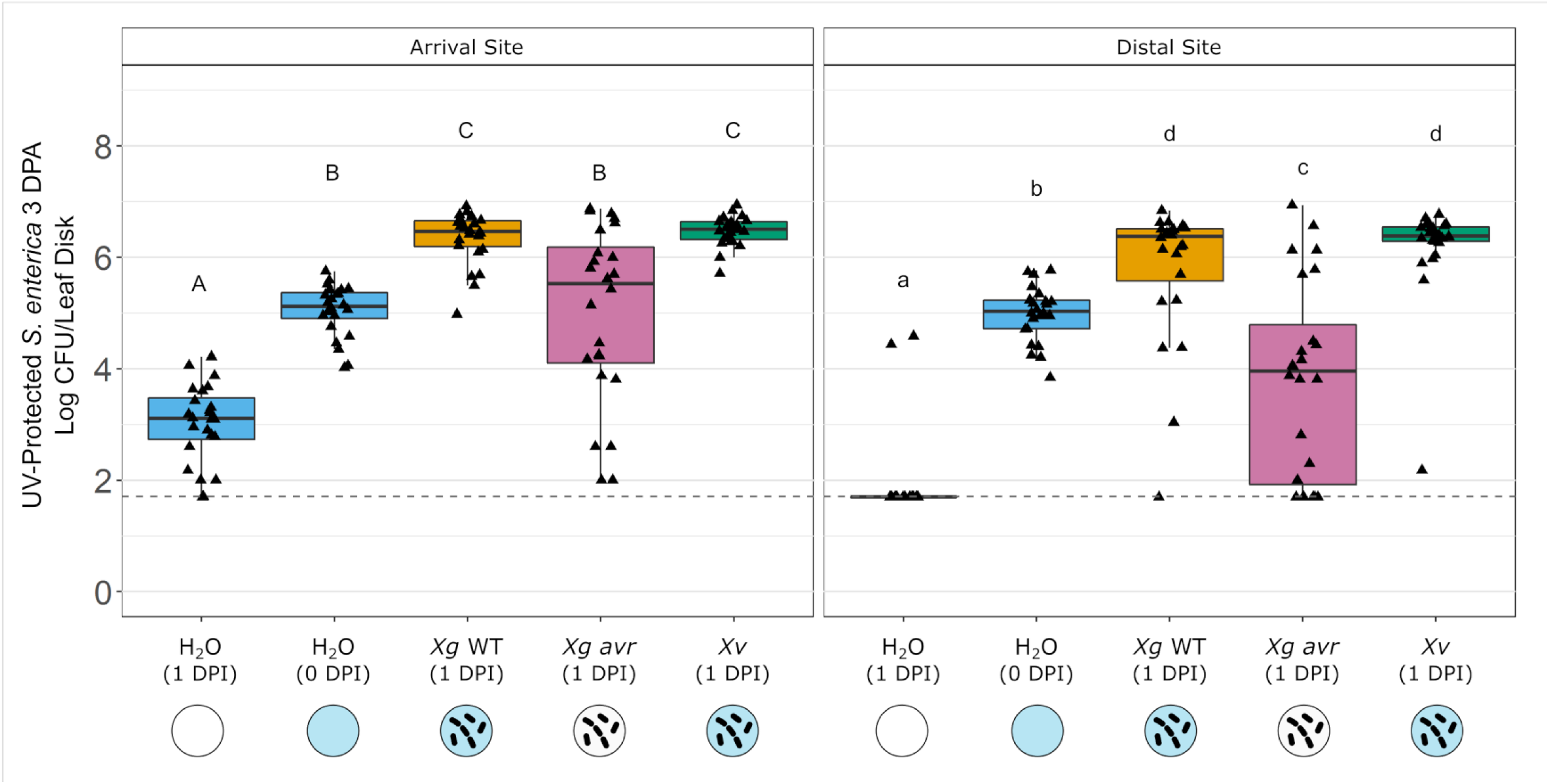
*Xanthomonas*-infected apoplast serves as a hospitable niche for *S. enterica* 3 days post- arrival. *S. enterica* (*Se*) was applied by droplet (10^6^ CFU/leaflet) to leaflets infiltrated with water (blue boxplots), *X. gardneri* WT (*Xg* WT; orange boxplots), *X. gardneri avrHah1^ΔDBD^* (*Xg avr;* pink boxplots), or *X. vesicatoria* (*Xv*; green boxplots) at indicated days post-infiltration (DPI). *Se* populations were sampled at 3 days post *Se*-arrival (DPA) from the arrival site and distal from the arrival site. All leaflets were UV irradiated prior to sampling. Uppercase letters above the boxplots denote significance among treatment groups within the arrival site based on an ANOVA and post-hoc Tukey test (*P* < 0.05). Lowercase letters denote significance among treatment groups within the distal site based on a Kruskal-Wallis and post-hoc Wilcox test (*P* < 0.05). Dotted line indicates limit of detection. Refer to Figure 1, panel C for circle symbol key describing apoplast states.

We also tested the hypothesis that *X. gardneri*-infected apoplasts support higher *S. enterica* growth when *S. enterica* cells are applied to the leaves at a 1000 times lower inoculum level (10^3^ CFU/droplet) relative to the previous experiments. Reducing the *S. enterica* inoculum dosage minimizes the possibility of overwhelming the plant immune response and more realistically models foliar contamination of crops with *S. enterica*. Arrival of *S. enterica* on healthy/dry leaves led to eradication of *S. enterica* populations in response to UV irradiation (Fig. 7). Meanwhile, when *S. enterica* was applied to leaves with an aqueous apoplast (healthy or *X. gardneri* WT-infected) it localized to UV-protected niches and migrated to distal tissue within 2.5 hours (Fig. 7). Additionally, the UV-protected populations increased between day 0 (2.5 HPA) and day 3 (3 DPA) (Fig. 7). *X. gardneri*-infected apoplasts supported the highest *S. enterica* growth, resulting in *S. enterica* populations that increased at least 3.8 logs within 3 days of arrival (Fig. 7). The higher apoplastic *S. enterica* populations observed in infected tissue substantiates the conclusion that prolonged survival in the apoplast by surface bacteria is supported by an aqueous environment resulting from phytobacterial infection, which is distinct from a healthy, watery apoplast.

**Figure 7:**
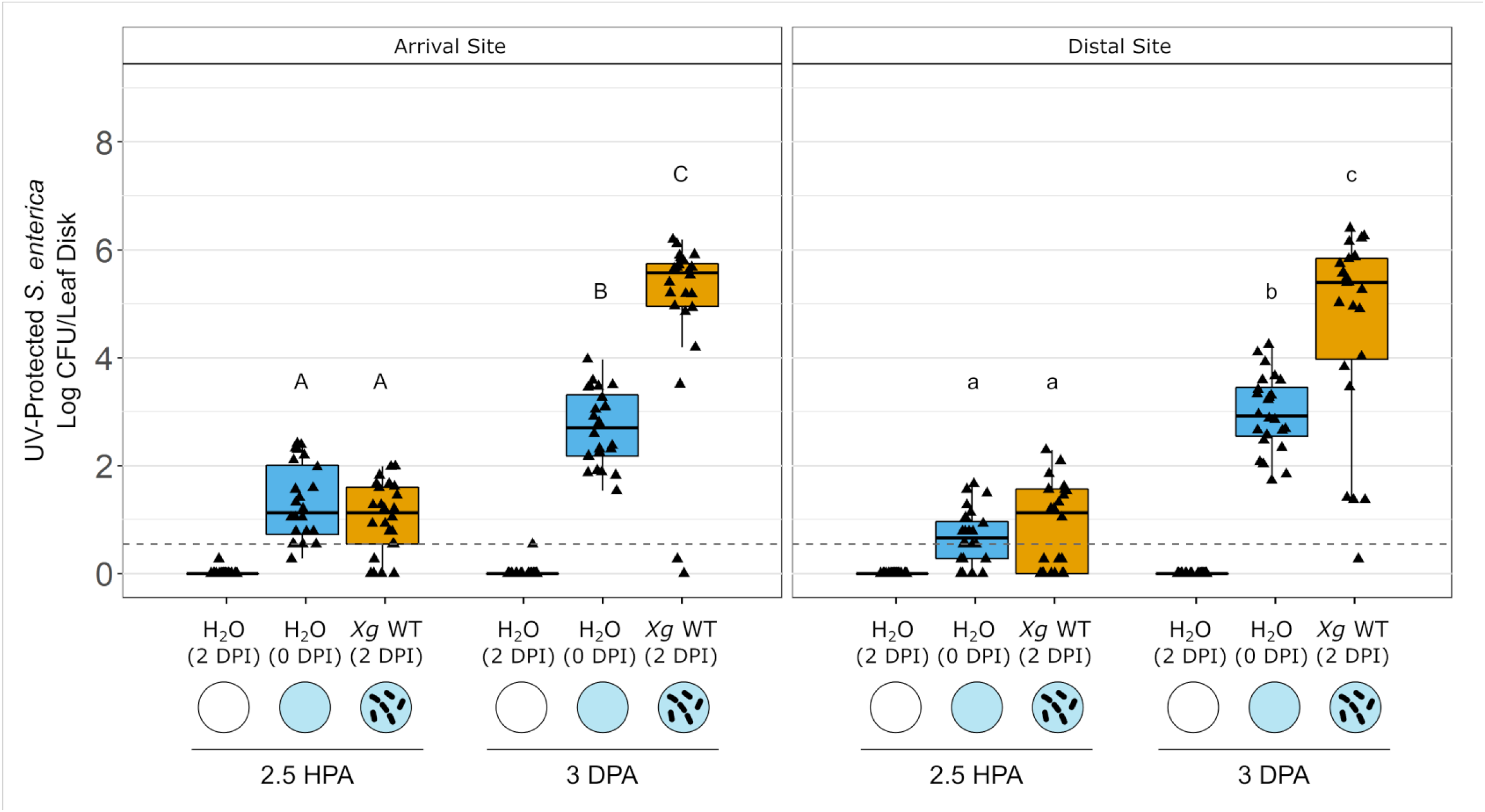
*X. gardneri*-infected apoplast enables replication of *S. enterica* within protected niches. *S. enterica* (*Se*) was applied by droplet (10^3^ CFU/leaflet) to leaflets infiltrated with water (blue boxplots), *X. gardneri* WT (*Xg* WT; orange boxplots), *X. gardneri avrHah1^ΔDBD^* (*Xg avr;* pink boxplots), or *X. vesicatoria* (*Xv*; green boxplots) at indicated days post-infiltration (DPI). *Se* populations were destructively sampled at 2.5 hours post *Se*-arrival (HPA) and 3 days post *Se*- arrival (DPA) at the droplet arrival site and distal from the arrival site. All leaflets were UV irradiated prior to sampling. Letters above the boxplots denote significance between the healthy/wet (H2O 0 DPI) and Xg treatment groups within the arrival (uppercase) and distal (lowercase) sites based on an ANOVA and post-hoc Tukey test (*P* < 0.05). Dotted line indicates limit of detection (LOD). Zero values were confirmed via LB enrichment. Points plotted at LOD/2 indicate absence of colonies via direct plating and a positive enrichment sample. Refer to Figure 1, panel C for circle symbol key describing apoplast states.

**Figure 8:**
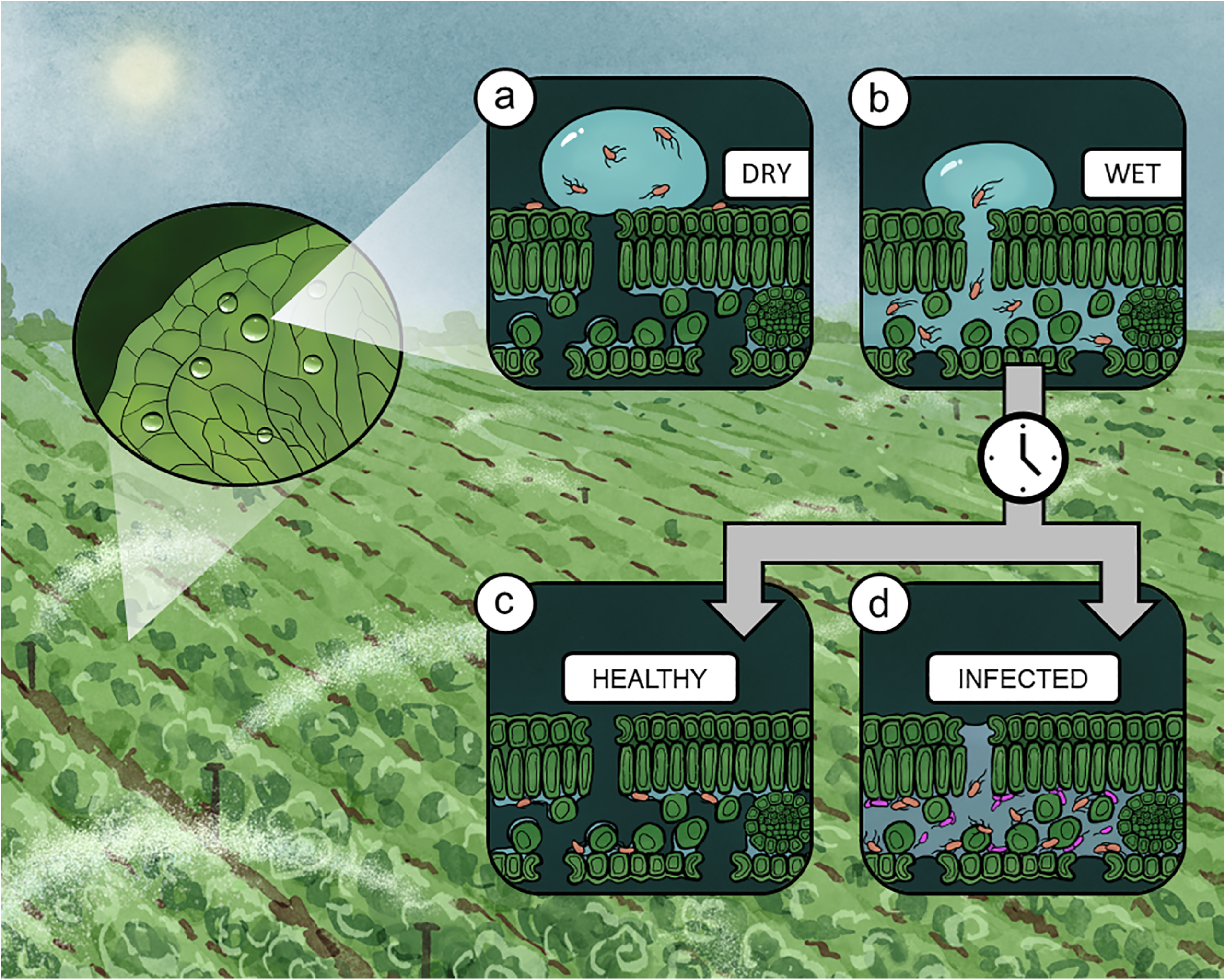
Model for *S. enterica* survival in a habitable apoplast. *S. enterica* can colonize crop plants via spray-irrigation of contaminated water. Macroscopically dry tissue (a) does not enable absorption of surface water, whereas tissue with an aqueous apoplast (b) facilitates the ingression of *S. enterica* into the apoplast. Healthy tissue that is abiotically water-soaked eventually dries out (c), creating an unfavorable niche for *S. enterica.* By contrast, an infected apoplast (d), which maintains water-soaking via phytopathogen effectors and is likely enriched with nutrients and other growth-promoting factors, facilitates *S. enterica* replication.

## DISCUSSION

In this study, we determined the fate of *S. enterica* following arrival on *Xanthomonas*- infected tissue, specifically regarding ingression into the apoplast and migration to distal sites. We hypothesized that the modifications to the host caused by *Xanthomonas* infection could facilitate ingression of *S. enterica* into the apoplast, where it could migrate to distal sites and thrive. *S. enterica* was applied to leaves as a droplet to model splash dispersal of bacteria in the field, which can occur during irrigation or precipitation events. Various host and microbial factors including the presence of water in the apoplast, flagellar function by *S. enterica,* and *avrHah1* expression by *X. gardneri* were manipulated to understand how alterations to the infection court, the apoplast, impact plant-associated surface bacteria.

Colonization of protected niches such as the apoplast is essential to the lifestyle of various plant-associated bacteria. To determine whether *S. enterica* could localize to protected niches of leaves, UV irradiation at 254 nm, which is an FDA-approved method for decontamination of food surfaces (31) and has been applied in previous plant bacterial endophyte studies (32, 33), was utilized to reduce surface *S. enterica* populations. The use of UV irradiation additionally models the UV exposure that plant-associated microbes are routinely exposed to, though it’s worth noting UV wavelengths less than 300 nm are typically absorbed by our atmosphere and do not reach Earth (34). Initially, we tested whether *S. enterica* cells that were infiltrated directly into the apoplast of leaves would be protected from UV irradiation applied to the adaxial surface, and we hypothesized that the palisade cells, upper epidermal cells, and cuticle would occlude UV penetration to the apoplastic *S. enterica*. We determined that *S. enterica* cells localized to the apoplast were more protected from UV compared to *S. enterica* on the surface, although UV irradiation reduced the apoplastic *S. enterica* populations a small but statistically significant amount (Fig. 2). Leaves must permit light to penetrate leaf tissue for photosynthesis and act as “light traps (35),” which may explain why a subset of apoplastic *S. enterica* cells were vulnerable to UV. The duration of UV exposure influenced the magnitude of *S. enterica* population reduction (data not shown), and the exposure level of 0.6 J/cm^2^ selected for these experiments resulted in >2.0 log reduction of surface *S. enterica* populations with minimal effects on the apoplastic populations. As for the surface-localized *S. enterica* cells, the small subpopulations that survived UV irradiation (Fig. S1) likely resisted sterilization by being canopied by leaf surface structures, such as trichomes, or wedged within the grooves between adjacent cells (36–38).

*S. enterica* successfully colonized UV-protected niches of abiotically water-soaked or *Xanthomonas*-infected tissue. We rationalized that *S. enterica* had accessed the leaf apoplast based on a few lines of evidence. First, the extent to which *S. enterica* was protected from UV irradiation following arrival on abiotically water-soaked or *X. gardneri-*infected apoplast was similar to cells infiltrated directly into the apoplast (Fig. 2). Second, *S. enterica* droplets were absorbed by abiotically water-soaked or infected tissue, as evident by rapid disappearance of the droplets in as little as 30 minutes. Healthy tissue is typically impermeable to spontaneous absorption of surface water because the surface tension of water is greater than the critical surface tension of the leaf surface (39). However, the relatively rapid disappearance of droplets into water-soaked leaves indicated that the wettability of the leaf surface had changed. Considering that the leaf tissue was deliberately not wounded, open stomata likely served as entry points for external water and suspended *S. enterica* cells to enter the apoplast. We theorize that an aqueous apoplast enabled a watery film to extend upwards through the stomatal pores, providing a continuous “watery highway” that could facilitate movement of *S. enterica* into the substomatal space. Lastly, *S. enterica* was consistently isolated at high levels from abiotically water-soaked (Fig. 2) and infected (Fig. 3) tissue ∼1 cm distal from the arrival site when the apoplast was water-soaked at *S. enterica* arrival, but not when it was dry. Schwartz et al. found that *X. gardneri*-infected leaves facilitated migration of surface *X. gardneri* cells to the apoplast (6), and here we have confirmed that other bacteria besides the phytopathogen *X. gardneri* could be transported by water-soaking. To verify that *S. enterica* colonization of distal tissue was not driven by migration across the surface but by migration through the apoplast, the adaxial surface of leaf tissue disks were pressed to agar plates following UV exposure. *S. enterica* was rarely isolated from the surface of leaves distal to the arrival site with a healthy/dry apoplasts (data not shown). Overall, *S. enterica* was found to relocate from the surface of leaves into the substomatal chamber and beyond. These observations posed the question of whether *S. enterica* motility is necessary for the redistribution of cells following *S. enterica* arrival.

Flagellar motility is key to survival on the leaf surface for recent immigrants (40), but we found that the relocation of *S. enterica* from the leaf surface to the aqueous apoplast can occur independent of functional flagella (Fig. 5), thus favoring a model of passive *S. enterica* relocation in the presence of a water-soaked apoplast. However, active motility was found to provide *S. enterica* with an advantage in colonizing sites beyond the initial arrive site at 2 days post-*X. gardneri* infection, but not earlier (Fig. 5). *In planta* microscopy of *Pseudomonas syringae* colonization has revealed that distinct bacterial subpopulations exist within infected tissue, suggesting that the apoplast is a heterogeneous environment (41). Later in *Xanthomonas* infection when more nutrients may be available in the apoplast, motile *S. enterica* cells may employ chemotaxis to colonize nutritional micro-oases within the apoplast. Therefore, as the *Xanthomonas* infection court dynamically changes over time, the importance of specific traits possessed by plant-associated microbes also changes.

In addition to determining how *S. enterica* redistribution is affected by one of its own genetic traits, we also examined the role of *X. gardneri* type-III effector AvrHah1 to identify mechanisms by which phytopathogen infection influences the fate of *S. enterica. AvrHah1* contributed to *S. enterica’s* ability to access the protective apoplast (Fig. 4) and migrate to distal tissue in the early stage of an ongoing *X. gardneri* infection (Fig. 5). AvrHah1 is the only *Xanthomonas* effector that has been directly linked to the water-soaking phenotype, despite water-soaking being a characteristic symptom of bacterial spot disease. Other Xanthomonads outside of the bacterial spot complex that infect diverse hosts, including leafy greens, cause water-soaking (42, 43), yet no other water-soaking effectors have been identified within the *Xanthomonas* genus. Among the various *Xanthomonas* species that cause bacterial spot disease, which includes *X. vesicatoria, X. euvesicatoria, and X. perforans, avrHah1* was originally specific to *X. gardneri* (44, 45). Recently, however, *X. perforans* strains carrying *avrHah1* have been identified in the southeastern United States (46). *X. vesicatoria,* which lacks *avrHah1,* induced visible water-soaking of tomato leaves as rapidly as *X. gardneri* and promoted protection and survival of *S. enterica* equal to or better than *X. gardneri.* These results contrast with previous studies documenting that *X. vesicatoria* infection does not promote *S. enterica* persistence on tomato leaf tissue following co-arrival of *X. vesicatoria* and *S. enterica* on healthy plants (15, 21). However, a key difference between those studies and the study presented here is the physiological state of the leaf when *S. enterica* arrives: healthy leaves versus infected leaves. Infected tissue can permit *S. enterica* to enter the apoplast, and therefore, the benefits conferred to *S. enterica* by *X. vesicatoria* infection may be conditional on *S. enterica* entry. Overall, the ability of *avrHah1-*lacking *Xanthomonas* strains to facilitate water-soaking mediated ingression of *S. enterica* into the apoplast and the observed delay, not elimination, of water-soaking in *X. gardneri avrHah1^ΔDBD^* suggest that other factors within the *Xanthomonas* genome independent of *avrHah1* contribute to creating an aqueous niche within the tomato leaf apoplast. Identifying these bacterial factors in future studies would improve our understanding of phytopathogen- mediated water-soaking.

Leaf surfaces are harsh environments that require plant-associated bacteria to flexibly adapt to extreme and dynamic conditions, including desiccation, heavy rainfall, UV irradiation, and sporadic nutrient availability (47–50). Many phytopathogenic bacteria have evolved to infect the apoplast of leaves where they can replicate to high population levels whilst exploiting host resources (51). The apoplast can be protective and provide bacteria with increased nutrient availability, however, this environment is exclusive to microbes that can successfully migrate there and tolerate the host immune response. Tomato leaves are relatively restrictive to *S. enterica* ingression (26, 27), yet we found that *S. enterica* can access an apoplast that is water- soaked either by *Xanthomonas* infection or abiotically. Furthermore, *Xanthomonas* infection resulted in higher *S. enterica* populations within UV-protected niches (Fig. 6) and enhanced *S. enterica* replication (Fig. 7) relative to healthy leaf controls. *S. enterica’s* capacity to exhibit near 4 log population growth within three days of *S. enterica’s* arrival on the surface of *X. gardneri*- infected leaves (Fig. 7) emphasizes the extent that *Xanthomonas* pathogens modify the apoplast. An infected tomato leaf apoplast relieves suppression of *S. enterica* replication, suggesting that *Xanthomonas* infection aids in overcoming the host immune response and/or liberates nutrients inaccessible to *S. enterica*. *S. enterica* is known to elicit the plant immune response (52) upon recognition of microbe-associated molecular markers such as flg22 (53, 54), although some evidence suggests that the host response provoked by *S. enterica* is relatively weak (55, 56). If modulation of the immune response by *Xanthomonas* does enhance *S. enterica* fitness in the apoplast, it is likely that this is not the only mechanism involved. The replication of *S. enterica* in an infected apoplast also suggests that nutrients have been made bioavailable by *Xanthomonas* infection. *Xanthomonas* spp. presumably liberate nutrients from leaf tissue, as supported by their own replication to populations above 10^8^ CFU/cm^2^ (21) and increased electrolyte leakage of tomato leaf tissue (15). One hypothesis is that an infected apoplast is enriched in carbohydrates, and *S. enterica* metabolizes these nutrients that it can’t access on its own. *S. enterica* can utilize many different sugars as sole carbon sources (57), and although only 10% of *Salmonella* species can metabolize sucrose (58), one of the most prevalent carbohydrates available in plant tissues (51), other sugars such as glucose and fructose (59, 60) could be available to *S. enterica*. Future studies will be informative in describing *avrHah1-*independent and -dependent mechanisms by which *Xanthomonas* releases host nutrients and identifying resources scavenged by *S. enterica* within infected leaves.

Overall, we have identified a common phytobacterial disease symptom of leaf pathogens, water-soaking, as a mechanism by which *S. enterica* can be passively relocated to the apoplast of leaves, resulting in access to protected niches including the substomatal chamber directly below the initial arrival site and distal sites of the apoplast. Our results show that *avrHah1* expression by *X. gardneri* influences the fate of *S. enterica,* and considering evidence that *avrHah1* is being transferred between *Xanthomonas* species, gene flow within a plant pathogen complex may have dire effects on the persistence of enteric human pathogens in crop plants that eventually will be consumed raw. Altogether, our findings illustrate a model for how enteric bacterial human pathogens are dispersed in agricultural fields via contaminated irrigation water and subsets of the bacterial population can relocate from the hostile leaf surface to a habitable niche within the apoplast (Fig. 7). As phytopathogen populations evolve and respond to global environmental changes, understanding their effects on the redistribution and survival of other plant-associated microbes will be critical.

## ACKNOWLEDGEMENTS

This material is based upon work supported by the National Science Foundation Graduate Research Fellowship under Grant No. DGE-1747503 and by USDA-HATCH grant no. WIS03022. Any opinions, findings, and conclusions or recommendations expressed in this material are those of the author(s) and do not necessarily reflect the views of the National Science Foundation. The funders had no role in study design, data collection, and interpretation, or the decision to submit the work for publication.

We are grateful to J. Jones and G. Minsavage for providing the *Xanthomonas gardneri* Δ*avrHah1* strain used in this work. Additionally, we thank C. Solís-Lemus and N. Keuler for providing guidance on statistical analysis.

## Author Contributions

MHD, KNC, and JDB developed the experiments. MHD, KNC, SCZ, and INM performed the experiments. MHD and JDB wrote the manuscript, and all authors reviewed the manuscript. MHD created the original illustrations.

