## Supplemental figures S1-S4, Table S1 for "*Xanthomonas* infection transforms the apoplast into an accessible and habitable niche for *Salmonella enterica*"

**SUPPLEMENTAL MATERIALS**

Figure S1:

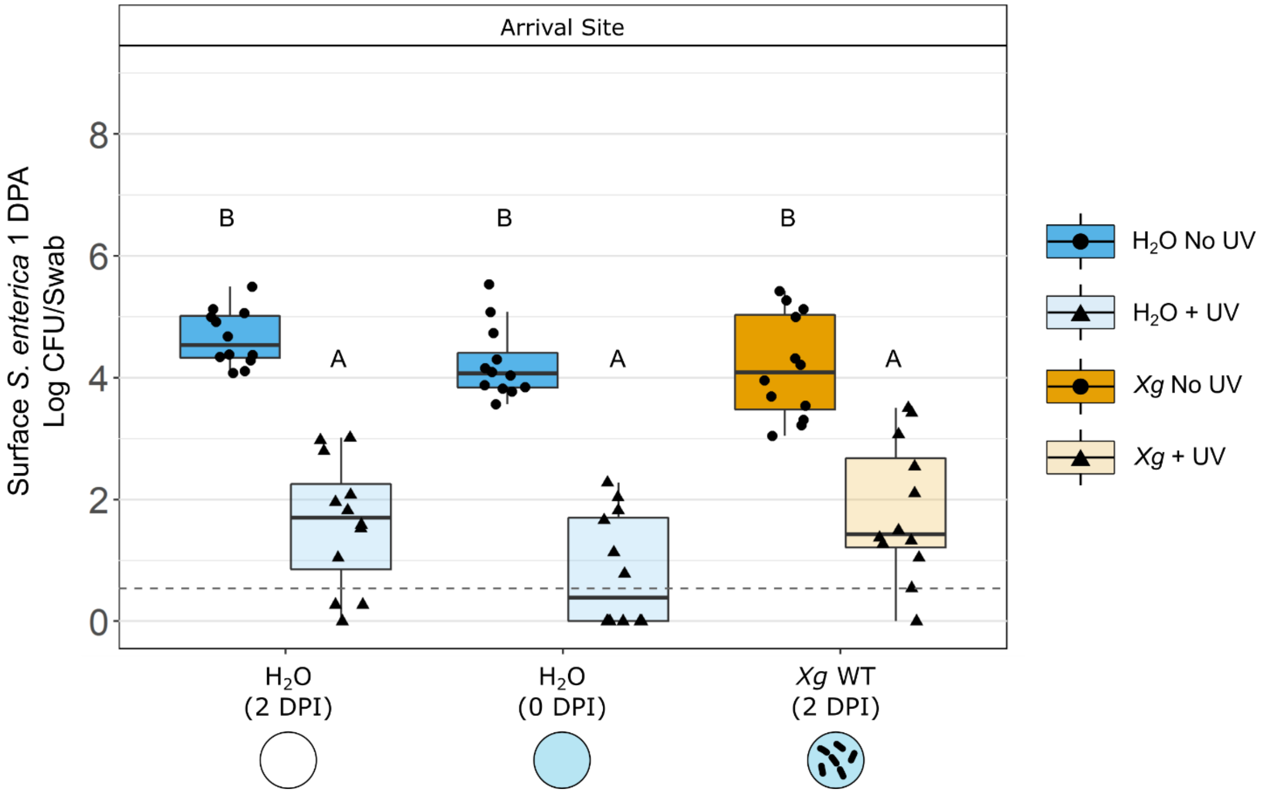

Figure S2:

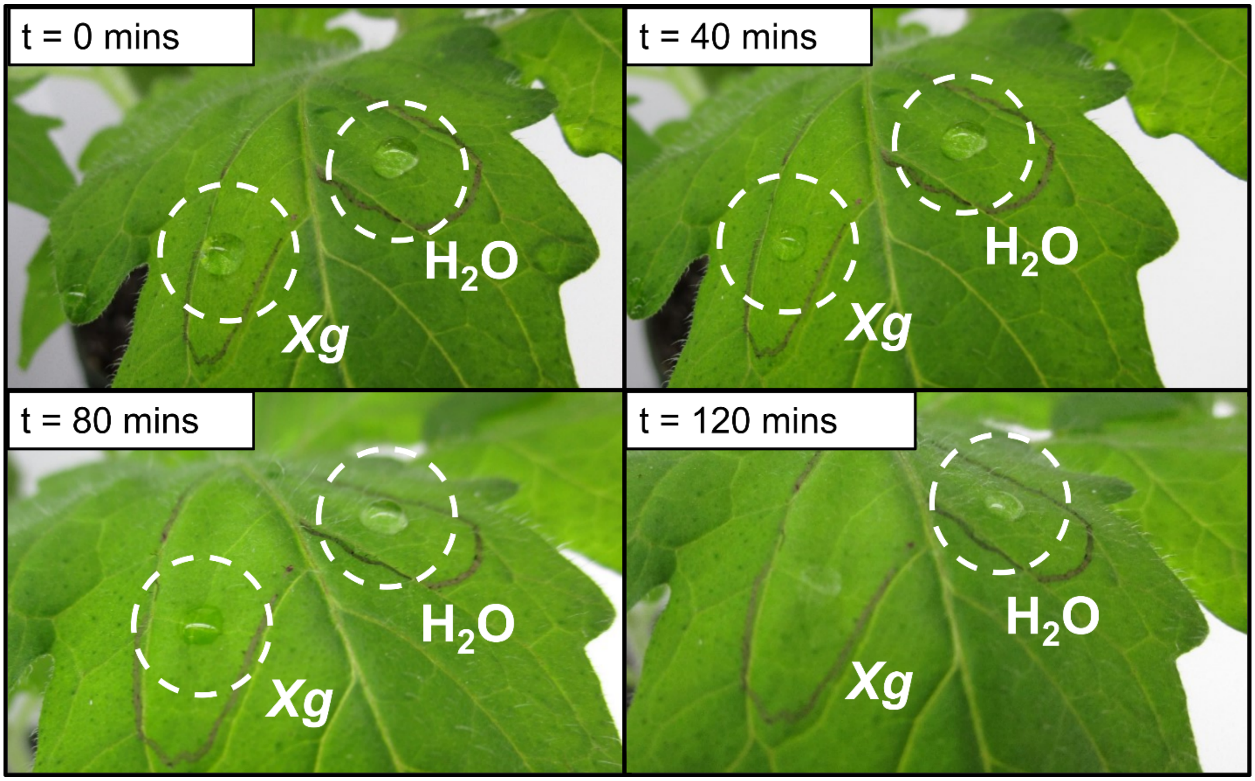


Figure S3:

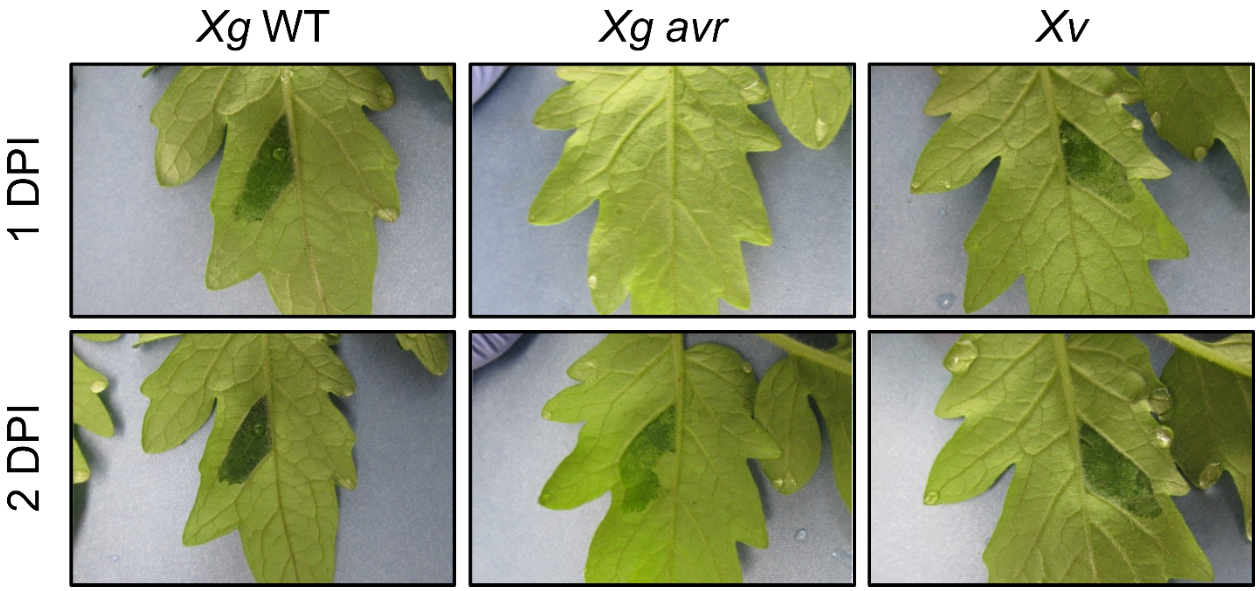

Figure S4:

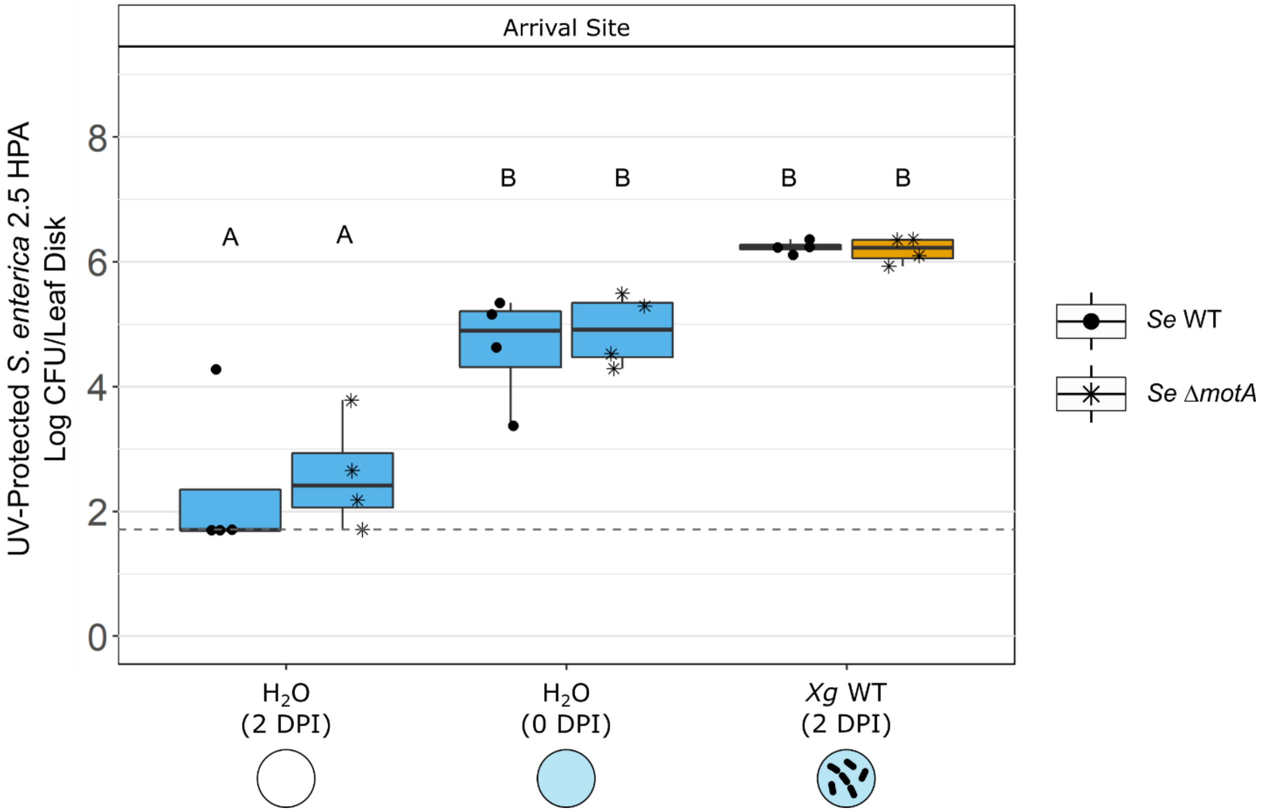


Table S1:

| Infiltration Treatment | Apoplast State at *Se* Droplet Arrival | Number of Droplets Completely Absorbed (n = 24) | Number of Droplets Partially/Not Absorbed (n = 24) | Incidence Rate of Complete Absorption (n = 24) |
|---|---|---|---|---|
| H2O 1 DPI | healthy/dry | 0 | 24 | 0% (A) |
| H2O 0 DPI | healthy/wet | 13 | 11 | 54% (B) |
| *X. gardneri* WT | infected/wet | 7 | 17 | 29% (B) |
| *X. gardneri* avrHah1$^{\Delta DBD}$ | infected/dry | 0 | 24 | 0% (A) |
| *X. vesicatoria* | infected/wet | 11 | 13 | 46% (B) |
